# The paternal and maternal genetic history of Vietnamese populations

**DOI:** 10.1101/719831

**Authors:** Enrico Macholdt, Leonardo Arias, Nguyen Thuy Duong, Nguyen Dang Ton, Nguyen Van Phong, Roland Schröder, Brigitte Pakendorf, Nong Van Hai, Mark Stoneking

## Abstract

Vietnam exhibits great cultural and linguistic diversity, yet the genetic history of Vietnamese populations remains poorly understood. Previous studies focused mostly on the majority Kinh group, and thus the genetic diversity of the many other groups has not yet been investigated. Here we analyze complete mtDNA genome sequences and 2.34 mB sequences of the male-specific portion of the Y chromosome from the Kinh and 16 minority populations, encompassing all five language families present in Vietnam. We find highly variable levels of diversity within and between groups that do not correlate with either geography or language family. In particular, the Mang and Sila have undergone recent, independent bottlenecks, while the majority group, Kinh, exhibits low levels of differentiation with other groups. The two Austronesian-speaking groups, Giarai and Ede, show a potential impact of matrilocality on their patterns of variation. Overall, we find that isolation, coupled with some limited contact involving some groups, has been the major factor influencing the genetic structure of Vietnamese populations, and that there is substantial genetic diversity that is not represented by the Kinh.

## Introduction

Southeast Asia (SEA) is a melting pot of ethnolinguistic diversity shaped by many demographic events, beginning with the initial arrival of anatomically modern humans at least 65 kya (1, 2), and including migrations accompanying the spread of agriculture, in particular rice and millet farming, the expansion of the Austronesian language family, and movements of Tai-Kadai and Hmong-Mien speakers (3). The languages spoken in SEA today belong to five language families: Austro-Asiatic (AA), Austronesian (AN), Hmong–Mien (HM), Sino– Tibetan (ST), and Tai–Kadai (TK). Geographically SEA is divided into two sub-regions, Island Southeast Asia (ISEA) and Mainland Southeast Asia (MSEA).

Vietnam is a multi-ethnic country that occupies a key position within MSEA and exhibits both geographic and ethnolinguistic diversity. The northern part of the country consists of highlands and the Red River delta; the central part also comprises highlands, while the southern part encompasses mostly coastal lowlands and the Mekong River delta. There are 54 official ethnicities in Vietnam that speak 109 different languages belonging to all of the five major language families present in SEA (4). Groups speaking AA languages are distributed throughout the country, while those speaking TK, HM, or ST languages are found in the north; AN-speaking groups are located in the central highlands. The AA language family is considered the oldest within the area; AA languages are scattered across MSEA and South Asia, and the location of the AA homeland is under debate (5). The AA languages are associated with a major occupation of MSEA after the introduction of agriculture (6).

AN speakers are found all over ISEA and Oceania, and trace at least a part of their ancestry to aboriginal Taiwanese AN, supporting a start of the AN expansion out of Taiwan about 4 kya (3, 7). The genetic composition of modern AN speakers in ISEA is heterogeneous; AN speakers in western Indonesia have substantial AA-related ancestry, caused most likely by a movement of AN speakers through MSEA mixing with AA speakers in Vietnam or peninsular Malaysia (3), while AN speakers in eastern Indonesia harbor both Papuan and AN-related ancestry (8). AN speakers in MSEA include the Cham, Chru, Raglai, Giarai, and Ede of central Vietnam, Cham of Cambodia, and Malay groups in Malaysia and Thailand. In contrast to the predominant patrilocal residence pattern of other groups, AN groups are thought to have an ancestral matrilocal residence pattern (9). The TK and HM languages likely originated in present-day southern China and then spread to Thailand, Laos, and Vietnam centuries ago (10, 11). Whether the current distribution of these languages and the farming culture across MSEA was a result of human migration events (demic diffusion) or happened without the major movement of people (cultural diffusion) is still highly disputed. MtDNA variation in Thailand supports a model of demic diffusion of TK speakers (12), while recent studies based on ancient DNA provide further evidence for Neolithic and Bronze age migrations from East Asia (13), and explain present-day SEA populations as the result of admixture of early mainland Hòabìnhian hunter-gatherers and several migrant groups from East Asia associated with speakers of the AN, AA and TK languages (14). ST languages are predicted to have diverged 5.9 kya in Northern China, and reached Vietnam around 0.5-0.15 kya (15).

While several genetic studies have focused on SEA, research on the ethnic groups in Vietnam remains rather limited (16–22). Most of these studies either focused solely on the majority group in Vietnam, the Kinh, as representative of the entire country, or are based on a restricted number of SNPs, microsatellites, or only partial sequencing of mtDNA. Because the Kinh comprise 86% of the population, a sampling of individuals from this group is a promising way to capture the main signal of genetic diversity in Vietnam. But the complicated history of SEA indicates that there might be hidden complexity and genetic structure in the minority populations. We have therefore initiated a comprehensive study of the genetic history of Vietnamese ethnolinguistic groups. Here, we analyze sequences of full mtDNA genomes and 2.3 million bases of the male-specific portion of the Y chromosome (MSY) of the Kinh and 16 minority groups, encompassing all five language families, to investigate their maternal and paternal genetic structure. We use the genetic results based on our extensive sampling to investigate whether the genetic composition of the Kinh is a valid representation of all populations living in Vietnam today, and we assess the impact of geographic, linguistic, and cultural factors (i.e., post-marital residence pattern) on the genetic structure of Vietnamese populations.

## Material and Methods

### Sample Information

We analyzed DNA from 600 male Vietnamese individuals (Supplemental Material Table S1) belonging to 17 ethnic groups that speak languages belonging to the five major language families in Vietnam. In detail the data set consists of two Austro-Asiatic (AA) speaking groups (Kinh and Mang), five Tai-Kadai (TK) speaking groups (Tay, Thai, Nung, Colao, and Lachi), two Austronesian (AN) speaking groups (Giarai and Ede), three Hmong-Mien (HM) speaking groups (Pathen, Hmong, and Dao), and five Sino-Tibetan (ST) speaking groups (Lahu, Hanhi, Phula, Lolo, and Sila). The average sampling locations per population are shown in Figure 1. Ethnic groups sampled for this project, name, language affiliation, and census size were based on *the General Statistics Office of Vietnam* (www.gso.gov.vn and the 2009 Vietnam Population and Housing census, accessed September 2018) and the Ethnologue (4). All sample donors gave written informed consent, and this research received ethical clearance from the Institutional Review Board of the Institute of Genome Research, Vietnam Academy of Science and Technology (No. 4-2015/NCHG-HDDD) and from the Ethics Commission of the University of Leipzig Medical Faculty.

**Figure 1.**
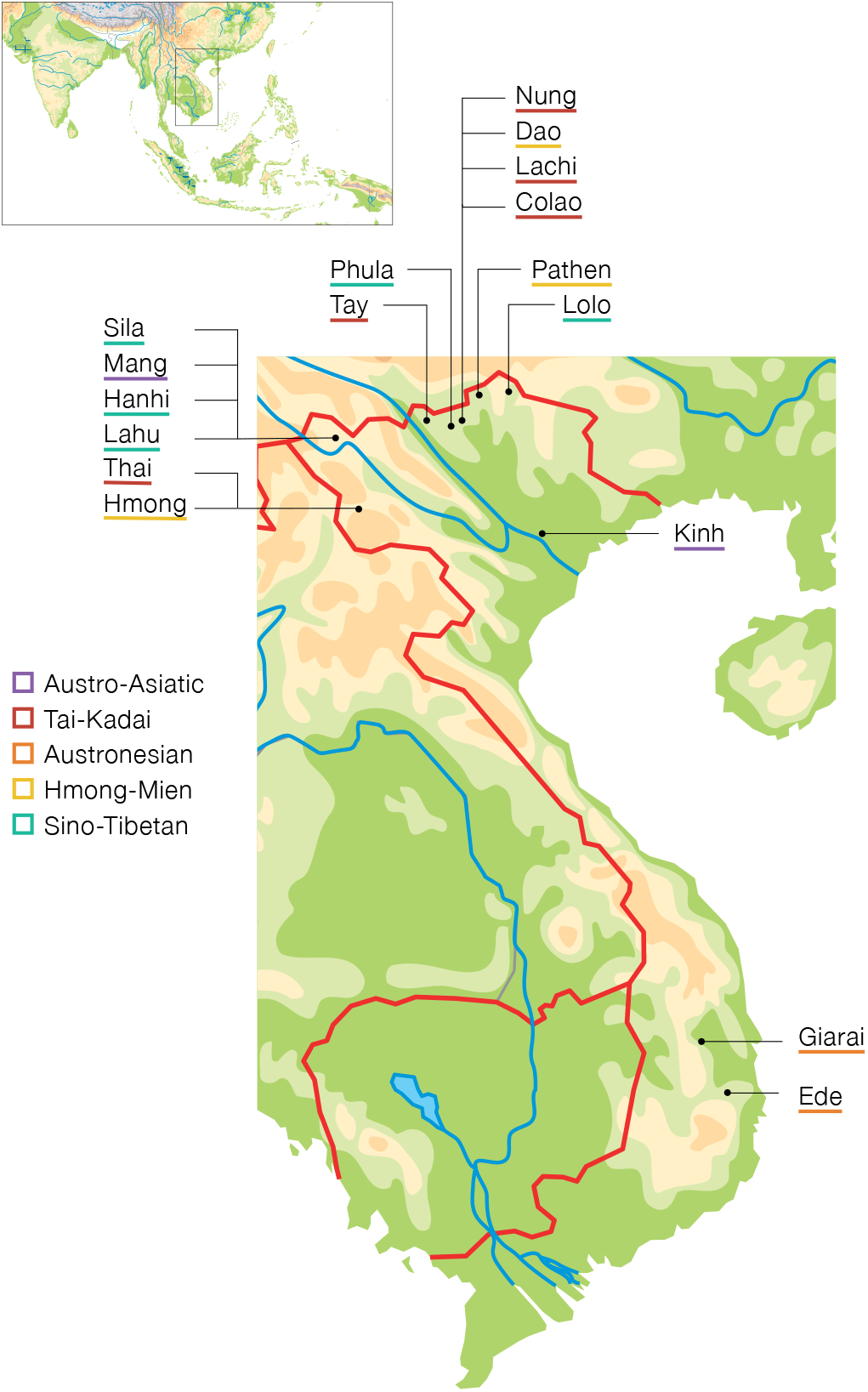
Map of sampling locations. Dots show average sampling locations per population. Population labels are color-coded by language family with Austro-Asiatic in purple, Tai-Kadai in red, Austronesian in orange, Hmong-Mien in yellow, and Sino-Tibetan in turquoise.

### MtDNA sequencing

DNA was extracted from blood using the Qiagen Tissue and Blood extraction kit. Double-stranded, double bar-coded Illumina sequencing libraries were constructed as described previously (23). The libraries were enriched for mtDNA sequences via in-solution capture (24). The mtDNA haplogroups were defined using haplogrep2 (ref. 25), and the phylogeny and terminology of Phylotree 17 (ref. 26) http://www.phylotree.org. For further experimental details see (27). The complete mtDNA dataset can be found at GenBank MH448947-MH449555.

### Y-chromosome sequencing

We enriched for ~2.3 million bases of the male-specific Y-chromosome (MSY) from the same libraries used for mtDNA capture-enrichment (27). The MSY sequence processing pipeline is described elsewhere (28). To increase the quality of the data set, we removed 41 positions with more than 16.6% missing information across the Vietnam MSY sequences and then imputed the remaining missing genotypes with BEAGLE 4.0 (ref. 29) using published reference sequences (Supplemental Material Table S2) from South Asia, East Asia, Southeast Asia, and Oceania (30–32). After making initial haplogroup calls (using the procedure described below), we then went back and added additional reference samples from haplogroups C2-M217 and N1-M2291 for imputation, as these were present in 28 and 22 individuals respectively in the Vietnamese sample set but not present in the initial set of reference sequences used for imputation. Further, a merged A00 sequence (30) was added as an outgroup. From the combined data set we additionally excluded 53 positions not covered by more than 75% of the samples. The aligned MSY reads are deposited in the European Nucleotide Archive (Project Accession ID available after peer review). Final SNP genotypes and their chromosomal positions on hg19 are provided in Supplemental Material Data 1 and Data 2. MSY haplogroups were called with yhaplo (33) using a stopping condition parameter ‘ancStopThresh’=10. Haplogroups were typed to the maximum depth possible given the phylogeny of ISOGG version 11.04 (http://www.isogg.org/) and the available genetic markers in our target region. Labels denoted with an asterisk in the text and figures are paragroups which do not include subgroups.

### Sequence analysis

For both markers, we calculated the mean number of pairwise differences by averaging over the sum of nucleotide differences for each pair of sequences within a population (R function: dist.dna package: ape) divided by the total number of pairs. The nucleotide diversity (*π*) and its variance were computed using the R function nuc.div (package:pegas). We calculated the number of unique haplotypes for each population and obtained the haplotype diversity (*H*) values using Arlequin (34) version 3.5.2.2. To visualize *π* and *H* we calculated the percentage difference from the mean for each population. Arlequin (34) version 3.5.2.2 was additionally used to calculate the pairwise genetic distances (Φ_ST_ distances) among the populations and the Analyses of Molecular Variance (AMOVA) for both markers. The p-values of the genetic distances were corrected for multiple testing by applying the Benjamini-Hochberg procedure. The Φ_ST_ distances were used to compute non-metric Multi-Dimensional Scaling (MDS) plots. We created two-dimensional projections (R function: isoMDS package: MASS) and calculated heatplots with 5 dimensions, showing per-dimension standardized values between 0 and 1. We calculated Mantel matrix correlation tests between genetic distances and great circle distances of the average geographical location per population using Pearson’s correlation with 10 000 times random resampling. The correspondence analyses were computed in R using the libraries ‘vegan’, ‘fields’ and ‘ca’. The haplotype sharing analysis was based on sequence haplotypes via string matching. We excluded Ns and indels for the mtDNA sequences; this step was not necessary for the MSY sequences because indels were not called and there were no Ns after imputation.

We performed mtDNA and MSY Bayesian analysis with BEAST 1.8 (ref. 35) The software jmodeltest2 (ref. 36) was used to determine that the HKY+I+G and GTR models were the best substitution models for the mtDNA and MSY sequences, respectively. We partitioned the mtDNA genomes into the coding and noncoding sections and applied previously published mutation rates (37) of 1.708 × 10 ^−8^ and 9.883 × 10 ^−8^ mutations/site/year, respectively. For all MSY analyses the MSY mutation rate of 0.871 × 10 ^−9^, based on an Icelandic pedigree (38), was applied. A Bayes Factor analysis including marginal likelihood estimations (39) was used to test different clock models. We chose the Bayesian Skyline piecewise linear tree prior for the dating and Bayesian Skyline generation. To ensure successful Bayesian estimation and to reach ESS values above 200, we combined multiple MCMC runs with 100 million steps using the BEAST logcombiner with a resampling up to approximately 40 000 trees.

We constructed MP trees for both markers and counted the mutations from the outgroup per sample. The mutation counts were used to compare the average distance of macro-haplogroups to the base of the trees as a measurement of branch length heterogeneity. We tested for significant differences in branch length distributions of major haplogroups with the Mann-Whitney-*U* test.

## 3. Results

### MSY sequences

We sequenced 2 346 049 bases of the MSY of 600 Vietnamese from 17 populations to a mean coverage of 30.2x (minimum: 5x, maximum: 72x). After filtering, there were 3932 SNP positions, including 2138 novel sites which have not been described previously (dbSNP Build 151, accessed 19^th^ September 2018 http://www.ncbi.nlm.nih.gov/SNP/). Fifty-seven different haplogroups were present in the 17 populations (Table S3). A detailed analysis of the phylogeography of the MSY dataset will be presented elsewhere (Nguyen et al. in preparation); the focus of the present study is the comparison of patterns of mtDNA and MSY variation in the sampled Vietnamese populations, and so we only briefly mention some interesting features about the MSY haplogroup distribution (Supplemental Materials Text 1, Figure S1, Table S3-4).

### Genetic diversity within populations

The nucleotide diversity (π) and haplotype diversity (*H*) for mtDNA and MSY sequences varied substantially among populations (Figure 2, Table S5). Kinh had high values of both π and *H* for both genetic markers, compared to the mean across populations. Sila, Lachi, and Mang had lower than average *H* values for both markers. The Mang MSY π-value was particularly low (5.91×10^−6^) compared to the mean (4.72×10^−5^), reflecting the unusual MSY haplogroup composition of this group, dominated by haplogroup O1b-B426, which had a frequency of 97%. The HM groups Pathen and Hmong had higher than average pi values for the MSY, which reflects the higher frequency of C and D haplogroup sequences in these two groups (Table S3). The two AN groups were notable in having substantially higher than average MSY *H* values but average or below average mtDNA *H* values. Overall, the variation in *H* and π values for both markers was not consistent within language families. We found high frequencies of both mtDNA and MSY haplotype sharing, up to 17 % (Figure 3), within all 17 populations. In total, 28.5 % of the mtDNA and 24.7 % of the MSY haplotypes were shared within at least one population. All groups shared mtDNA types within the population, and only the Giarai lacked any shared MSY haplotypes within the population. The highest frequencies of within-population MSY haplotype sharing were present among the Mang (0.15), Sila (0.17), and Colao (0.14), in keeping with their very low *H* values (Figure 2).

**Figure 2.**
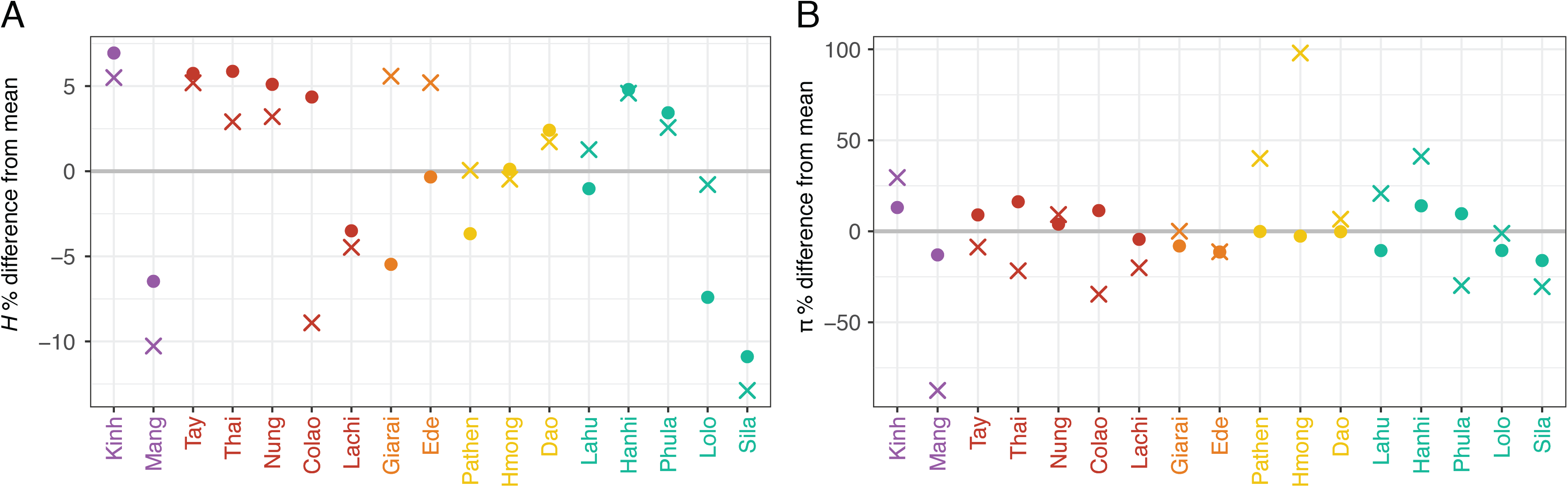
Diversity statistics, shown as the percent difference from the mean of: A: the haplotype diversity (*H)* and B: the nucleotide diversity (π). Crosses and dots denote the MSY and mtDNA values, respectively. Population labels are color-coded by language family with Austro-Asiatic in purple, Tai-Kadai in red, Austronesian in orange, Hmong-Mien in yellow, and Sino-Tibetan in turquoise. The grey line shows the mean across populations.

**Figure 3.**
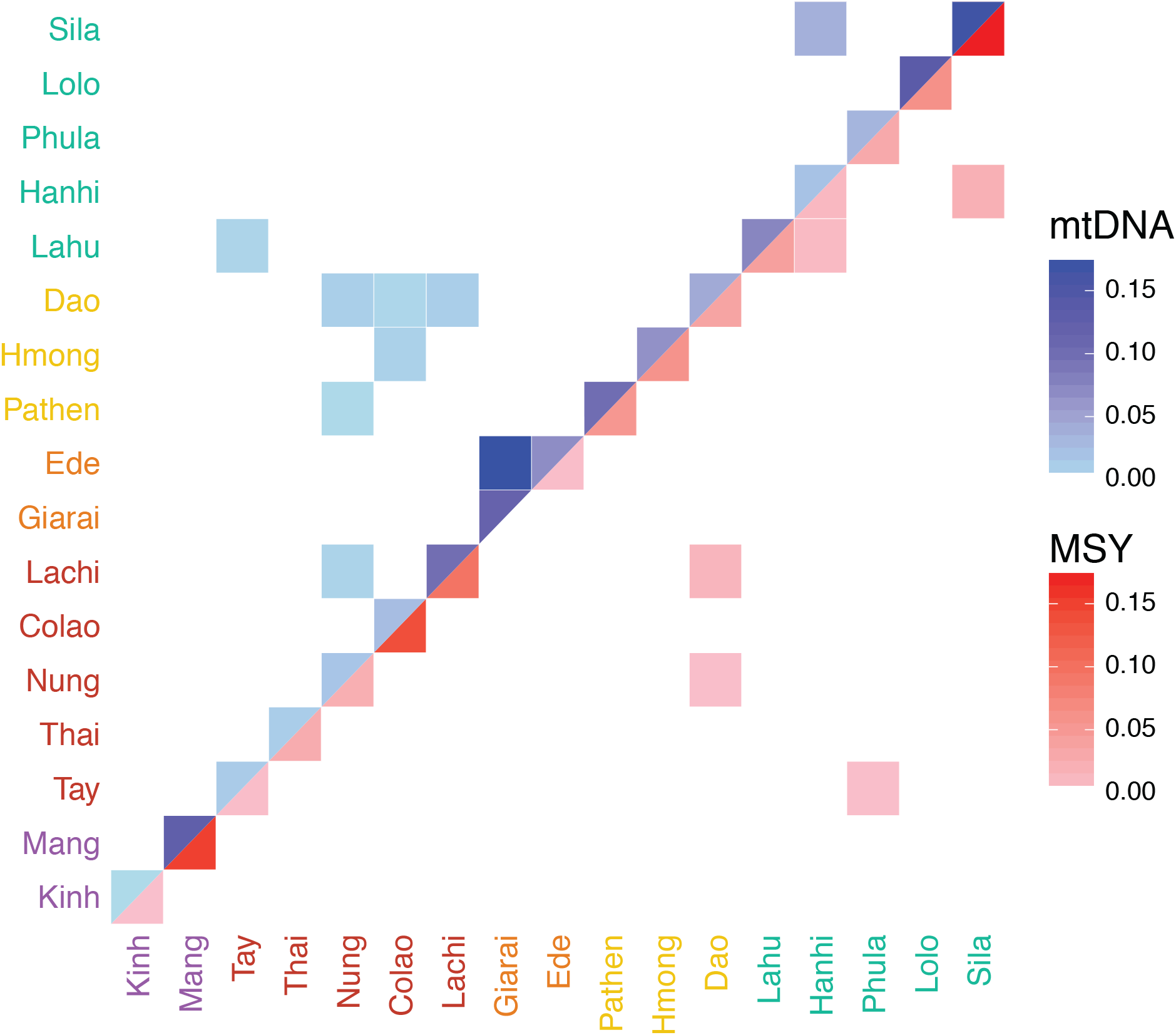
Frequency of shared haplotypes between populations. mtDNA (upper triangle) and MSY (lower triangle) shared haplotype frequencies are represented by the blue and red color scale, respectively. White squares indicate no sharing. Population labels are color-coded by language family with Austro-Asiatic in purple, Tai-Kadai in red, Austronesian in orange, Hmong-Mien in yellow, and Sino-Tibetan in turquoise.

### Population relationships

We examined haplotype sharing between populations as an indication of recent genetic contact or shared ancestry (Figure 3). In general, there were more occurrences of sharing between populations in the mtDNA sequences compared to the MSY sequences, but only 5.5 % of the mtDNA and 2.8 % of the MSY haplotypes were shared between populations. The two AN populations (Giarai and Ede) had the highest frequency of mtDNA haplotype sharing between them (0.16), which was even higher than their within-population sharing (0.10 and 0.07). However, in notable contrast to the high degree of mtDNA sharing, there was no MSY haplotype sharing between the AN groups. The three HM populations each shared mtDNA haplotypes with some TK populations, but not with other populations of their own language family. The Dao (HM) shared MSY haplotypes with the TK groups Nung and Lachi; they pare located close to one another and also shared mtDNA haplotypes. All other occurrences of MSY haplotype sharing involved populations in close geographical proximity, namely Sila-Hanhi, Lahu-Hanhi, and Phula-Tay. To visualize the relationships among Vietnamese populations, we generated MDS plots (Figure 4) from the matrices of pairwise Φ_ST_ distances (Supplemental Material Figure S2) for both mtDNA and the MSY. Sila and Mang exhibited large distances to all populations, which explains their position in the periphery of the MDS plots (Fig 4A and 4B). Additionally, Phula (ST) stands out in the MSY MDS plot. Giarai and Ede showed large Φ_ST_ distances for mtDNA but not for the MSY (Figure 4), and larger Φ_ST_ distances to most ST and HM populations than to Kinh (AA) and TK (except Lachi) groups. The Kinh group showed overall low genetic distances with other groups (Supplemental Material Figure S2) and a central position in both MDS plots (Figure 4). Because the rather high stress values of the two-dimensional MDS plots (Figure 4A and 4B) indicated potentially more complex structure, we calculated a five-dimensional MDS and depicted the results in a heat plot (Supplemental Material Figure S3A and S3B). The Kinh, Thai, and Tay remained centrally-located across all 5 dimensions for both markers (Supplemental Material Figure S3A and S3B), while the Mang remain an outlier in most dimensions in the MSY plot (Supplemental Material Figure S3B).

**Figure 4.**
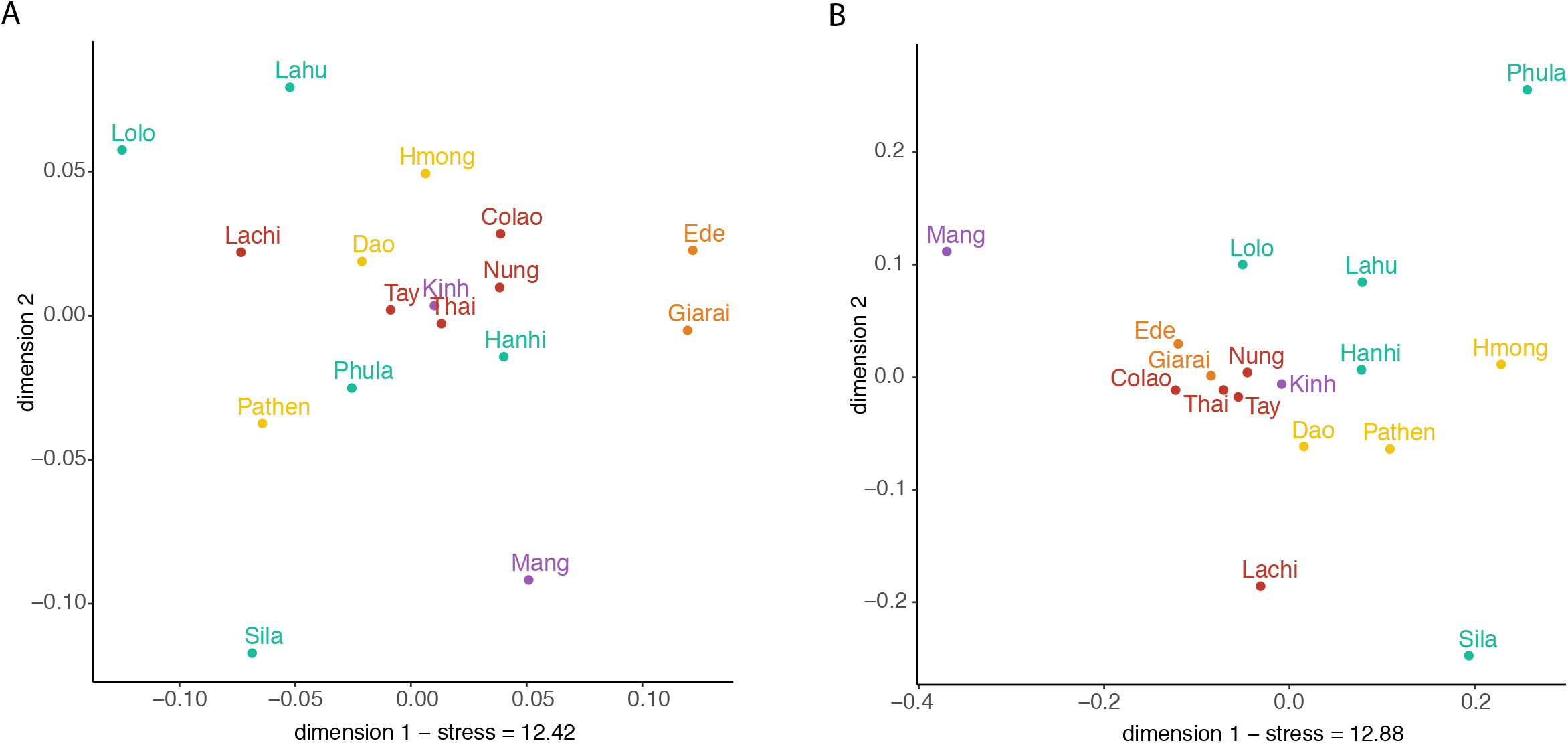
A: mtDNA and B: MSY MDS plots based on Φ_ST_ distances. Stress values are in percent. Population labels are color-coded by language family with Austro-Asiatic in purple, Tai-Kadai in red, Austronesian in orange, Hmong-Mien in yellow, and Sino-Tibetan in turquoise.

We additionally explored population relationships based on haplogroup frequencies via a correspondence analysis (CA) (Supplemental Material Figure S4). The results were similar to the MDS results for the AN groups, in that the Giarai and Ede were outliers for mtDNA but not for the MSY. Their mtDNA separation, in the first dimension of the plot, was driven mainly by the high frequencies of haplogroup M71 + C151T (37-42%), as well as by other exclusive haplogroups such as M68a1a, F1a4a, M21b, M24b, M68a1a, M73b, M74b1, M7b1a1f, and R9b1a1a. The second dimension separated the Mang (haplogroup A, D4, M71) and Pathen (A14, F1d,F2a). For the MSY, the high frequencies of F-M89 separated Phula (74%) and Lahu (32%) from the rest in the first dimension, while the second dimension spreads the populations between Mang and Hmong (Figure S4).

### Factors influencing the genetic structure

To test for correspondence between linguistic affiliation or geographic location with genetic structure, we analyzed three groupings (linguistic affiliation and two levels of geographical proximity) via AMOVA (Table 1). With respect to geography, we grouped populations on a broad scale by regions (political units), and on a finer scale by their origin in the same or neighboring districts. All three tested grouping patterns (language family, district, and region) revealed that ~90% of the total mtDNA variation and ~77% of the total MSY variation is explained by the differences within populations. Although the among-group component was significant for language family (1.8%) and districts (2.6%) for the mtDNA sequences, and for language family (4.4%) for the MSY sequences, in all of these cases the within-group component was considerably larger, indicating that differences between populations assigned to the same group were bigger than differences between populations assigned to different groups.

To further assess the impact of geography on the genetic structure of Vietnamese populations, we tested for correlations between the geographic and genetic distances for both mtDNA and MSY sequences (Table 2). This analysis was carried out for the entire data set, for a subset excluding the Kinh, and for a subset excluding the Ede and Giarai. We excluded the Kinh to control for the influence of their geographically widespread distribution and potentially mixed gene pool, as they are the majority group in Vietnam. Giarai and Ede, the only two groups from the Central Highlands of Vietnam, were excluded to test for a potential bias caused by their unique geographic position, as including these groups results in a bimodal distribution of geographical distances. We found a significant correlation between the geographic distance and the mtDNA genetic distance matrices when analyzing the entire data set and the population subset excluding the Kinh (Table 2). However, the correlation between mtDNA distances and geography became non-significant when excluding the two Central Highlands groups (Ede and Giarai), suggesting that their large geographic distance (Figure 1) and high mtDNA genetic distances (Figure S2) from the other groups was driving the significant correlation. Furthermore, no significant correlation was detected for any comparisons of MSY genetic and geographic distances. However, there was a significant correlation between the genetic distance matrices of the two uniparental markers (Table 2) when the Kinh were included.

We additionally carried out maximum parsimony and Bayesian analyses of the MSY sequences and tested for branch length heterogeneity; the results of these analyses (which indicate substantial branch length heterogeneity in the MSY tree, but not the mtDNA tree) are discussed in Supplemental Material Text 2 and 3.

## 4. Discussion

Previous genetic studies of Vietnam have focused largely on the Kinh majority as representative of the country (14, 16-18, 20, 21, 32, 40, 41). In contrast, we have investigated the patterns of genetic variation in a large sample of ethnolinguistic groups from Vietnam that speak languages encompassing all of the five language families present in the country. We found varying levels of genetic diversity within and among groups, not all of which is represented in the Kinh. For example, the high genetic diversity observed within the Kinh and several TK speaking groups differed substantially from the much lower levels of diversity found in small populations like the Sila and the Mang (Figure 2). The predominant sharing of haplotypes within populations (Figure 3), as well as this reduced diversity (Figure 2), suggests that many of the Vietnamese groups are relatively isolated from one another. The higher number of population pairs which share mtDNA haplotypes, compared to MSY sharing, likely reflects recent contact and is in accordance with the expectation of more female exchange due to patrilocality in all non-AN groups. However, lower levels of MSY sharing could also reflect the greater resolution of MSY haplotypes based on 2.3 Mb of sequence, compared to the 16.5 Kb mtDNA genome sequences. Considering the difference in mutation rates for the different molecules (37, 38), a new mutation is expected in an MSY haplotype of our target size every 489.4 years, and in the whole mtDNA genome every 3624 years; thus, new mutations erase MSY sharing faster than mtDNA sharing. The proportion of MSY haplotypes which are shared within Vietnamese populations (24.2%) is much higher than observed in previous studies of the same MSY regions in other populations (7.1% for Thai/Lao populations (42), 13.7% for northwest Amazonian populations (28), and 6.9% for Angolan populations (43) (Supplemental Material Figure S7).

There was no correspondence between the genetic structure and geography, as indicated by the absence of a significant correlation between genetic and geographic distances (Table 2), and the lack of any significant influence of geographical clusterings in the AMOVA (Table 1). While the mtDNA genetic structure is slightly influenced by geography, a significant correlation between mtDNA genetic distances and geographic distances disappears when the AN groups are excluded (Table 2). Because there are other factors which differ between AN groups and non-AN groups, such as the post-marital residence pattern (discussed in more detail in Supplemental Material Text 4), which might influence the genetic structure, geography is not necessarily directly related to genetic structure. We also did not find any evidence for an association between genetic structure and language family affiliation (Table 1). A consistent finding across Vietnamese populations was a higher female than male effective population size (Supplemental Materials Text 2, Figure S5), and more genetic structure on the MSY (MSY F_ST_ = 23% vs. mtDNA F_ST_ =10%) (Table 1). These are common patterns in human populations (41, 44), and likely reflect a predominant patrilocal residence pattern and higher levels of female migration (45–47), and a greater variance in reproductive success for males than for females (46, 48). Strikingly, the Pathen (HM) mtDNA and MSY effective population sizes were about the same. Why this is the case is not known, but we speculate that this could reflect greater homogeneity in male reproductive success for the Pathen, compared to other Vietnamese groups.

Overall, it appears that the genetic structure of Vietnam has been primarily influenced by two main factors. The first is isolation and genetic drift, leading to high levels of genetic differentiation between groups and variable levels of genetic diversity within groups. Second, there has been limited contact between some groups, leading to some haplotype sharing. The levels of genetic differentiation among groups of 10% based on mtDNA and 23% based on the MSY (Table 1) are similar to what was found previously for isolated populations from Northwestern Amazonia (13% mtDNA and 27% MSY; (28)) and higher than those found for Thai populations (8.5% mtDNA and 11% MSY; (42)). We also found particularly low levels of diversity in some specific groups, like the Mang and Sila (Figure 2, Supplemental Material Table S5). The low levels of haplotype sharing for both markers (Figure 3) are further evidence of isolation and limited contact between geographically close populations. The observed level of mtDNA haplotype sharing between Vietnamese groups (5.5%) is lower than that observed in most other studies of complete mtDNA genome sequences (Supplemental Material Figure S6), while the MSY haplotype sharing between Vietnamese groups (2.2%) is also lower than what was observed in studies that sequenced the same regions of the MSY (Supplemental Material Figure S7).

In addition to the general aspects of Vietnamese genetic diversity discussed above, our results provide some insights into the genetic profile and history of specific groups. These are discussed in detail in Supplemental Text Materials Text 4, and include: the higher male than female isolation of HM groups; recent expansions and diversification of TK groups, along with some contact between them and HM groups; the impact of matrilocality on patterns of genetic variation in the AN groups; evidence for the probable incorporation of other groups into the Kinh during their initial spread; and the pronounced bottleneck (especially in the MSY sequences) in the Mang and Sila.

## Conclusion

The sequencing of 2.34 Mb of the male-specific Y-chromosome and the complete mtDNA genome of 17 different ethnic groups (encompassing all five language families) enabled us to carry out the first comprehensive analysis of the genetic diversity within Vietnam. Overall, isolation leading to genetic drift has had an important impact on Vietnamese groups, with recent contact limited to some TK and HM groups. There are several differences between the maternal and paternal genetic history of some populations; in particular, a matrilocal versus patrilocal residence pattern appears to be one of the major drivers of differences between the MSY and mtDNA signals in AN vs. other groups. There is also a profound impact of genetic drift for the Mang and Sila, especially in the MSY lineages, suggesting male-specific bottlenecks or founder events. And although we find genetic evidences for a central position of the Kinh as the majority group within the country, there is substantial genetic diversity in the other ethnic groups that is not represented in the Kinh. Genetic studies of the remaining ethnic groups in Vietnam, and expansion of the genetic data to include genome-wide variation, will provide further insights into the genetic history of this key region of MSEA.

Supplementary information is available at European Journal of Human Genetics’ website.

## Acknowledgments

We thank Alexander Hübner for computational advice, and Wibhu Kutanan and Dang Liu for helpful discussions. This research was supported by funds from the Max Planck Society and the Ministry of Science and Technology, Vietnam (Grant DTDL.CN-05/15). BP is grateful to the LABEX ASLAN (ANR-10-LABX-0081) of Université de Lyon for its financial support within the program "Investissements d’Avenir" (ANR-11-IDEX-0007) of the French government operated by the National Research Agency (ANR).

## Table captions

Table 1 – AMOVA results. Populations included in each group for each classification are indicated below the table. Significance *, <0.05; ***, <0.001

Table 2 – Correlation coefficients obtained in the Mantel correlation tests. Significance * p < 0.05.

## Supplementary captions and titles

Table S1 – Sample information. MtDNA haplogroup labels are based on Phylotree 17, while MSY haplogroups are designated in short form by the diagnostic SNP and in long form by the haplogroup name based on ISOGG version 11.04.

Table S2 – MSY reference data set used for imputation.

Table S3– MSY haplogroup frequencies per population. “MSY CA cluster” indicates how haplogroups were combined for the correspondence analysis.

Table S4 – mtDNA haplogroup frequencies per population. “MtDNA CA cluster” indicates how haplogroups were combined for the correspondence analysis.

Table S5 – Diversity statistics for a: MSY and b: mtDNA sequences per population including sample size, number of haplotypes, nucleotide diversity (*π*) and standard error (*π* SE), haplotype diversity (H), and the mean number of pairwise differences (MPD).

Table S6 – Mann-Whitney U tests for comparisons of the number of mutations to the outgroup for pairs of haplogroups.

Table S7 – Bayes factors for comparing a strict clock model and an uncorrelated relaxed lognormal clock model.

Figure S1 – Macro-haplogroup frequencies per population for mtDNA (left) and MSY (right), in percent.

Figure S2 – Pairwise Φ_ST_ distances between populations, with mtDNA and MSY represented by the blue and red color scale, respectively. Distances that are not significantly different from zero are marked with a tilde symbol (p> 0.05). Population labels are color-coded by language family with Austro-Asiatic in purple, Tai-Kadai in red, Austronesian in orange, Hmong-Mien in yellow, and Sino-Tibetan in turquoise.

Figure S3 – A: mtDNA and B: MSY multidimensional heatplot based on Φ_ST_ distances with 5 dimensions and values for each dimension scaled to be between 0 (blue) and 1 (red). Population labels are color-coded by language family with Austro-Asiatic in purple, Tai-Kadai in red, Austronesian in orange, Hmong-Mien in yellow, and Sino-Tibetan in turquoise.

Figure S4 – Correspondence analysis plots for A: mtDNA and B: MSY sequences, based on haplogroup frequencies. Grey vectors and labels show haplogroup projections. Population labels are color-coded by language family with Austro-Asiatic in purple, Tai-Kadai in red, Austronesian in orange, Hmong-Mien in yellow, and Sino-Tibetan in turquoise.

Figure S5 – MSY and mtDNA Bayesian Skyline Plot (BSP) for all populations.

Figure S6 – Proportion of shared mtDNA haplotypes within and between populations in Vietnam vs. other studies of complete mtDNA genome sequences. 1: Barbieri, C. et al. 2014 Am. J. Phys. Anthropol.; 2: Barbieri, C. et al. 2011 Mol. Biol. Evol.; 3: Mizuno F. et al. 2014 J. Hum. Genet.; 4: Arias, L et al. 2018 Am. J. Phys. Anthropol.; 5: Duggan, A. T. et al. 2014 Am. J. Hum. Genet.; 6: Delfin, F. S. et al. 2014 Eur. J. Hum. Genet.; 7: Duggan, A. T. et al. 2013 PLOS ONE. 8: Gunnarsdóttir, E. D. et al. 2011 Nat. Commun.; 9: Barbieri, C. et al. 2013 Am. J. Hum. Genet.; 10: Ko, A.-M.-S. et al. 2014 Am. J. Hum. Genet.; Vietnam: this study.

Figure S7 – Proportion of shared MSY haplotypes within and between populations in Vietnam vs. other studies of the same MSY region. 1: Oliveria et al. 2019 Hum. Genet.; 2: Arias et al. 2018 Am. J. Phys. Anthropol.; 3: Kutanan et al. 2019 Mol. Biol. Evol.; Vietnam: this study.

Figure S8 – Maximum parsimony tree of Vietnamese MSY sequences labeled with macro-haplogroups.

Figure S9 – Comparison of the number of mutations to A00 for selected MSY macro-haplogroups

Figure S10 – Maximum parsimony tree of Vietnamese mtDNA sequences with haplogroup labels.

Figure S11 – Comparison of the number of mutations to the RSRS for selected mtDNA macro-haplogroups

Figure S12 –MSY Bayesian Skyline plot (BSP) for the Giarai, Ede, and Kinh.

Figure S13 – BSP of major MSY macro-haplogroups, including sub-groups with a sample size > 20: C2e-F2613, F-M89, N1-M2291, O1a-M119, O1b-M268, and O2a-M324.

Figure S14 – BSP of MSY macro-haplogroup O1b-M268 sub-haplogroups. The asterisk denotes a haplogroup without subgroups.

Figure S15 – BSP of MSY macro-haplogroup O2a-M324 and sub-haplogroups.

Figure S16 – Bayesian tree of MSY haplogroup O1b-B426 (O1b1a1a1b1a) including the cluster of Mang individuals. The TMRCA point estimate and 95% HPD interval are shown in red.

Figure S17 – Bayesian tree of MSY haplogroup O1b-F1759 (O1b1a2a1) including the cluster of Sila individuals. The TMRCA point estimate and 95% HPD interval are shown in red.

