## SupplementMaterial Text for "The paternal and maternal genetic history of Vietnamese populations"

Supplemental Material: Text

1. **MSY haplogroup distribution**

The number of haplogroups per population ranged from two to 21; no haplogroup was found in all populations. The haplogroup distribution within the two AA populations (Mang and Kinh) varied considerably. In the Kinh, 21 different haplogroups were found, with O1b-F2758* having the highest frequency (22 %; Supplemental Material Table S3). Conversely, there were just two haplogroups in the Mang (AA), with O1b-B426 having a frequency of 97% (Supplemental Material Table S3). This is a sub-haplogroup of O1b-M1283*, which was present in all language families, and overall was the most frequent haplogroup (15%). O2a-F8 was also found in all language families, but its subgroups were more specific for groups of populations: we found O2a-F155 and O2a-F317 only in the Tay (TK), but O2a-F2137, O2a-F46*, and O2a-F4110 in Dao (HM) and in Lahu and Hanhi (ST).

All TK groups had high frequencies of O-M175 sub-haplogroups (overall 95%; Supplemental Material Table S3) ranging from 88% in Colao to 100% in Lachi. In contrast, the HM groups were characterized by high frequencies of haplogroups C and D. In particular, 63% of all Hmong individuals belonged to sub-haplogroups of haplogroups C-M130 or D-M174.

The AN groups (Ede and Giarai) were the only groups with haplogroups L-M27, R2-M479 and subgroups of R1-M173. Furthermore, Ede and Giarai individuals predominantly carried O1b-M1283*, the most common haplogroup in this study, at frequencies of 58% and 59% respectively. The Ede and Giarai also stood out with respect to mtDNA haplogroups; in particular, they had high frequencies of M71 + C151T (37% and 42%, respectively), which was absent elsewhere (Supplemental Material Table S4).

The ST groups were characterized by various haplogroups found at high frequency in just a few groups. For example, the Sila had a high frequency (83%) of O1b-F1759, which otherwise was found in only one other ST group (Hanhi, 9%) and in the Kinh (2%). The Phula had the highest frequency of F-M89 (47%), followed by Lahu (32%) and Hanhi (15%). This haplogroup was also found in low frequencies (< 7%) in two additional ST groups, and in Dao (HM), Tay and Thai (TK). The Lolo had a high frequency (69%) of haplogroup O1b-Z24091, which otherwise was found in the TK-speaking Lachi (22%) and at frequencies of <10% in a few other groups (Supplemental Material Table S3).

2. **Maximum Parsimony and Bayesian analysis**

The MSY Maximum Parsimony tree (Supplemental Material Figure S8) shows an overview of the paternal lineages, in which haplogroup O1-M1354 and subgroups thereof predominated. The average number of mutations to the outgroup per macro-haplogroup (Supplemental Material Figure S9) showed high levels of heterogeneity, and there were highly significant differences (even after Bonferroni correction for multiple comparisons) between branch length distributions for 23 of the 28 pairwise comparisons between macro-haplogroups (Supplemental Material Table S6). In contrast, the mtDNA tree (Supplemental Material Figure S10) shows less branch length heterogeneity (Supplemental Material Figure S11). Only six significant differences in branch length distributions between major mtDNA haplogroups were detected (Supplemental Material Table S6), of which five differentiated the longer branches of mtDNA macro-haplogroup F from all others.

In light of this branch length heterogeneity, in the Bayesian analysis we tested for different clock models (strict clock vs. an uncorrelated lognormal relaxed clock) by calculating the Bayes factor support from the Marginal Likelihood estimations (see Methods and Supplemental Material Table S7). The results indicated very strong or decisive support for the relaxed clock model for the complete MSY data set, and for MSY subsets for 12 of the 17 populations. In contrast, the complete mtDNA sequence set, each MSY haplogroup subset, and the MSY population subsets from Lachi, Lahu, Phula, Tay, and Thai showed either weak support for a relaxed clock or weak support for the strict clock. To avoid overfitting, we only selected the more complex relaxed clock model for sequence sets with at least strong support for the relaxed clock.

We constructed BSPs to study the changes in the mtDNA and MSY population sizes over time for all 17 ethnic groups (Supplemental Material Figure S5). The general pattern revealed a lower effective population size of the MSY than of mtDNA, although with some exceptions. An expansion of the female effective population size was present across the data set starting between 50 and 40 kya. Most population BSPs curves exhibited a very recent decrease within the last few thousand years. This signal could be the sign of a population contraction or a known methodological artifact due to potential substructure within populations (1). Some groups deviated from this general pattern: the MSY BSP of Ede (AN) and Kinh (AA) showed an increase of the effective male population size during the last 2.5 kya, while the Giarai (AN) showed a similar increase but at an earlier time, around 7.5 kya (Supplemental Material Figure S12). In comparison, the mtDNA estimates of population sizes over time indicated that, in addition to the 50-40 kya expansion, Tay and Kinh exhibit an additional expansion starting 15 kya (Supplemental Material Figure S5). Also, while most populations exhibited consistently lower MSY than mtDNA-based effective population sizes, for the Pathen (HM) the mtDNA and MSY effective population sizes were about the same.

In order to gain further insights into the differences in population history indicated by the BSPs for the different groups, we also constructed BSPs by MSY haplogroups (Supplemental Material Figure S13). We found that within O1b-M268 only the sub-haplogroup O1b-M1283* (O1b1a1a1b) exhibited two events of population increase (Supplemental Material Figure S14) starting at 2 and 5 kya. The BSP of macro-haplogroup O2a-M324 indicated specificity within its sub-haplogroups as well. The detailed analysis reveals a differentiation between the signals of O2a-F2588 (O2a2a1) and O2a-P164 (O2a2b), which suggests trends for population increases starting at 2 kya and 6 kya, respectively (Supplemental Material Figure S15). Both increases were succeeded by a population contraction. The population size history of MSY macro-haplogroup F-M89 (Supplemental Material Figure S13) dated back to only 2.5 kya and is characterized by a very steep increase starting at 2 kya. As the populations exhibiting strong signals of recent population increase in the MSY (Kinh Giarai, and Ede) lacked haplogroup F-M89, their signal of recent population expansion instead appeared to be driven by subhaplogroups of O1b-M268 and O2a-M324 (Supplemental Materials Figures S13, S14, S15).

In keeping with the high MSY haplotype sharing and low diversity found in the Mang, 36 of the 37 MSY sequences formed a monophyletic cluster with a common ancestor dated to 1.9 kya (1.2 – 2.5 kya 95% HPD Interval; Supplemental Materials Figure S16). The MSY BSP for the Mang correspondingly indicated a trend for a decreasing population size and a very short history. The male population size trajectory for the Sila (ST) indicated a similar trend. The majority (83 %) of the Sila individuals carried MSY haplogroup O1b-F1759 (O1b1a2a1), which coalesced very recently at ~1.6 kya (Supplemental Materials Figure S17).

3. **Branch length heterogeneity**

We found significant branch length heterogeneity for the MSY, but not for mtDNA, which adds to the complexity of the genetic structure between the two uniparentally inherited markers. The fact that the samples were all collected and processed in the same manner excludes potential branch length biases caused by batch effects; furthermore, sequence coverage across samples was high and relatively even, excluding an effect from the imputation. The comprehensive reference data set used for the imputation further avoids a branch length reduction via underrepresentation of singletons. In addition, our results are in keeping with previous reports of branch length differences in the MSY phylogeny (2-4). The signal we found in our dataset suggests that branch length heterogeneity may be associated with population expansions or effective population size differences. The shortest branches on average were present in MSY haplogroups D-M174 and F-M89 (Supplemental Material Figure S9). For haplogroup F-M89* which has been previously reported in India, Pakistan, Sri Lanka, Nepal, Borneo, Java, (5) and Iran, (6), we found highest frequencies among the three ST-speaking groups (Phula, Lahu, and Hanhi, Table S3), a lower effective starting population size compared to all other macro-haplogroups (Supplemental Material Figure S13), and expansion beginning only ~ 2 kya; all other major haplogroups had about one magnitude higher effective population size and older expansions (Supplemental Material Figure S13). The sample size for macro haplogroup D-M174 was too low to estimate a reliable BSP. The reason for this branch length heterogeneity is unknown; it has been previously suggested that an older paternal age for Bantu farmers compared to Khoisan foragers, and hence a higher mutation rate, could contribute to the rate heterogeneity among African MSY haplogroups (2). Further studies are needed to determine if paternal age differences could also play a role in the MSY rate heterogeneity in Vietnam.

4. **The genetic profile of Vietnamese populations**

In this note we provide more details concerning the genetic profile and history of some of the Vietnamese populations.

***The Hmong***

The Hmong-Mien (HM) language family contains 32 languages and is one of the major language families spoken in EA and SEA. The Proto-Hmong-Mien language is dated back to 2.5 kya or 4.3 kya (7, 8). The maternal history of HM groups points towards an origin in Southern China (9), and a spread further south and towards Vietnam 0.2-0.3 kya (10). The mtDNA also indicates some contact with the north among the Hmong groups (which are grouped as Miao in China) (9). A recent forensic Y-STR study found that Hmong people from the Chinese Yunnan region are most closely related to other populations from southern China, in particular from the Hunan region, the proposed homeland of the Dao people (11).

The MSY haplogroups we found in the Hmong (HM) belong predominantly to early diverging macro-haplogroups C-M130 and D-M174 (Supplemental Material Table S3), which explains their very high diversity at the nucleotide level (Figure 2, Supplemental Materials Table S5). The Pathen (HM) also exhibited higher frequencies of C-M130 and elevated MSY nucleotide diversity compared to other groups (Figure 2, Supplemental Materials Table S3 and Table S5); however, the other HM group, the Dao, did not have such high frequencies of these haplogroups. As the HM speaking populations are thought to be indigenous to Southern China (12), and to have migrated from there to Vietnam, our results suggest that sequences belonging to C-M130 and D-M174 are likely to be characteristic of HM groups and have been maintained at relatively high frequencies over time, perhaps due to little incorporation of male lineages from other groups into the Hmong. The mtDNA haplogroup distribution indicates a less specific pattern (Supplemental Materials Table S4, Figure S1), potentially caused by a higher rate of mtDNA contact with other groups. Notably, HM groups share mtDNA haplotypes with TK groups (Figure 3), suggesting extensive recent contact between HM and TK groups, as found in a previous study of Vietnamese mtDNA variation (13).

***TK groups***

The TK language family is the most numerous in terms of speakers among Vietnamese minorities, ranging from ~1.5 million for the Tay and Thai down to the low thousands (~ 2600 Colao) or even below (687 in the Pu Péo, not part of this data set) (the 2009 Vietnam Population and Housing census). While the language family is thought to have originated in Southern China, it expanded within the last 2 ky and diversified to a total of 91 different languages spoken all across MSEA and China, and is particularly predominant in Thailand and Laos (14). Our data set included groups speaking Central Tai languages: the Tay and the Nung (Zhuang in China); Southwestern Tai languages: the Thai; and Kadai languages: the Colao and the Lachi. The latter two have a closer genetic connection to the Laotian TK speakers and to the HM groups (Wibhu Kutanan, personal communication). There are some similarities in the diversity values of TK groups from the Tai branch, as the Tay, Thai and Nung have *H* values much above average for both markers, and MSY π values that are also above average (Figure 2, Supplemental Material Table S5). However, there is no uniform pattern for either the entire language family or the separate Tai and Kadai branches, as there are several exceptions: the Nung have the only above average MSY *π* value; the Colao have low MSY *H* and π values; while the Lachi have low *H* and π values for both markers (Figure 2, Supplemental Material Table S5). The TK groups Tay, Thai, Nung, and Colao tend to be clustered together towards the center of the MDS and CA plots, while the Lachi are more of an outlier (Figure 4, Supplemental Materials Figure S4). The BSP plots of population size change over time (Supplemental Materials Figure S5) also suggest different population histories for different TK groups. The Tay BSP is characterized by an mtDNA expansion (15 kya) and trend for MSY expansion (5kya), while the BSP plots for the other TK groups do not indicate such expansions. The fairly high levels of diversity within TK groups and limited differentiation between groups are in keeping with a recent expansion and diversification of TK groups. However, the previously noted mtDNA and MSY sharing between HM and TK groups has also had an impact on the history of some TK groups, suggesting a more complex story than a simple expansion into MSEA.

***The* *Giarai and Ede***

The Giarai and Ede are the only two AN-speaking populations sampled for this study, and they are also the only two sampled groups from the Central Highlands (Figure 1). They speak closely related languages belonging to the Malayo-Polynesian branch of Austronesian languages (10, 15), which are widespread in ISEA but rare in MSEA. Their genetic history has not been previously studied; however, they are probably related to the Cham, another AN group situated in the Vietnamese central highlands that has been studied (16, 17). The mtDNA lineages of the Cham include haplogroups (B, R9, M7, M17, M21, M22, M50, M51, M73, M77, N21, R22, and R23) which link them to MSEA, and specifically to the Kinh, rather than to ISEA populations. Therefore, it was hypothesized that the spread of the AN languages in MSEA was mediated by cultural diffusion, which was facilitated by the importance of central Vietnam as a trade network hub (16). The Giarai and Ede show high frequencies of the mtDNA haplogroup M71 + C151T which is specific to Vietnam (18). It is a subhaplogroup of M71 which is found across MSEA but is absent in Taiwanese AN speakers (18). The MSY haplotypes of the Cham (O-M95* and P-P27.1) have been associated with both MSEA and ISEA populations through STR haplotype sharing and haplogroup network analysis (17). Specifically, the C-M216, F-M213*, and K-P131* haplogroups indicate recent gene flow from ISEA, while O-M7, O-M88, O-M134, O-P191*, and O-P200* indicate a closer connection to MSEA populations (17). The suggested recent MSY geneflow from ISEA indicates that the spread of AN languages in MSEA was not exclusively culturally driven. We found a majority of the Giarai and Ede sequences in O-M1283 (Supplemental Material Table S3), which is a descendant of O-M95 and was found across all language families in our study. Even though the different depths of haplogroup typing complicates the comparison, there are some similarities in the haplogroups or subgroups found in the Giarai and Ede with the Cham, namely B, R9, M7, M21, M73, N21 for the mtDNA, and C-M216 and the aforementioned O-M95 subgroups for the MSY.

The Giarai and Ede are unusual in having much higher than average values of MSY haplotype diversity, whereas their mtDNA haplotype diversity values are average (Ede) or even below average (Giarai) (Figure 2). The pronounced differences between the two AN groups and the other Vietnamese groups may reflect differences in residence pattern, as all 15 non-AN groups included in this study practice patrilocality, while the AN groups practice matrilocality (19). Patrilocality has been suggested to influence patterns of genetic diversity (20), and a previous study of hill tribes of northern Thailand (21) found contrasting patterns for matrilocality that are similar to the Giarai and Ede vs. other Vietnamese groups. However, studies of other populations have failed to show a matching pattern of diversity and post-marital residence (22-24). This is not surprising; as discussed previously (25) there can be flexibility in post-marital residence pattern, and many other factors (subsistence, isolation, contact, etc.) can influence patterns of diversity as well. Nonetheless, the predominant patrilocal background in Vietnam is probably behind the substantially higher proportion of variation among populations for the MSY than for mtDNA (Table 1). Because Giarai and Ede are the only two matrilocal groups in the data set, dispersing men and Y-chromosomes could account for their comparably high levels of within-population MSY diversity (Figure 2), more central position in the MDS plot for the MSY than for mtDNA (Figure 4), and more similar haplogroup profiles to other groups for the MSY than for mtDNA (Supplemental Material Figure S4). However, we did not observe any MSY haplotype sharing between Giarai and Ede, or between them and any other group. This may reflect the limited sampling of Central Highland populations in the current study. It is also not clear if the sharing of mtDNA haplotypes between Giarai and Ede is an indication of a specific shared history or a general feature of Central Highland groups; further sampling and analysis of other AN and non-AN groups in the Central Highlands would be desirable.

***The Kinh***

The Kinh speak an AA language of the Vietic branch (Ethnologue) and are the majority group in Vietnam (accounting for 86% of the population). They are widely distributed across the country, although our samples are only from Hanoi and the surrounding area. The origin of the Kinh ethnic group is thought to go back to ancient settlers in the Phong Chau region in Nothern Vietnam (10, 19).

It is likely that the large census size and wide distribution of the Kinh reflect both expansion and incorporation of other groups. This is reflected in their high levels of genetic diversity for both markers (Figure 2) and their central position in all comparative analyses (Figure 4, Supplemental Material Figure S4). Additionally, mtDNA and MSY genetic distances were correlated when including the Kinh (Table 2), suggesting that contact between other groups and the Kinh has been an important factor. The BSPs for the Kinh indicate an increase within the last 2.5 ky for the MSY (Supplemental Material Figure S12) and an older increase for the mtDNA (Supplemental Material Figure S5). Therefore, the rise of this AA group in Vietnam might be linked to the spread of the Dong Son culture 2.6 kya (26), which coincides with the expansion of the MSY effective population size. As the Kinh are the most widespread (and prestigious) ethnic group in the country, contact with many other ethnic groups would be expected, and the absorption of individuals from other ethnicities would increase the diversity of the Kinh genepool and reduce their genetic distances to other groups. However, if contact or adoption of individuals to the more prestigious Kinh was very recent, we would expect some degree of sequence sharing, which we did not find. We would expect the MSY haplotype based on the callable region in this study and a mutation rate of 0.871 x 10 ^-9^ (ref. 27) to receive a new mutation on average every 489.4 years or 16 generations, and so recent contact (during the past few hundred years) should result in MSY haplotype sharing. Thus, it is likely that the high genetic diversity and central position in the MDS and CA plots of the Kinh reflects amalgamation and incorporation of other groups during their initial spread, rather than more recent contact.

***The Mang***

The Mang language is an AA language of the Northern Mon-Khmer branch with in total only 4205 speakers (14). This language is spoken both in Vietnam (census 3900) and by about 500 individuals in the Yunnan province of China. There is a report that the Mang in China may have migrated from Vietnam (28).

In our data the Mang show extraordinarily low MSY diversity (Figure 2, Supplemental Material Table S5), with all but one individual belonging to haplogroup O1b-B426 (Supplemental Materials Table S3, Figure S16). These results agree with a previous analysis of Chinese Mang MSY diversity based on 12 SNPs and 8 microsatellite loci (29). While the Mang mtDNA genepool also exhibits low genetic diversity (Figure 2, Supplemental Material Table S5) and high genetic differentiation (Figure 4), they do have several mtDNA haplogroups (Supplemental Material Table S4) and the BSP for mtDNA is comparable to those for the other Vietnamese groups (Supplemental Material Figure S5). It appears that the Mang have lived for an extended time in isolation, and/or undergone a pronounced male-specific bottleneck, with a reduction to one primary MSY lineage about 1.9 kya (Supplemental Materials Figure S16). Because of the shared low MSY diversity with the Mang in China (29), the divergence between these two groups must have happened more recently than 1.9 kya.

***The Sila***

The Sila are an ST speaking group with Chinese origins and linguistic similarities to the Lolo and Hanhi (14, 19). It is proposed that Sila individuals migrated to Vietnam in the 1830s from the Phongsali province, one of the areas in Laos where 3150 of the in total 3860 Sila individuals (14) are living today (10, 30).

The Sila exhibit the lowest *H* values for both mtDNA and the MSY (Figure 2, Supplemental Material Table S5) and high genetic distances to other groups (Figure 4, Supplemental Material Figure S2). As with the Mang, the majority (83%) of the MSY sequences belong to just one haplogroup, O1b-F1759 (Supplemental Material Table S3). Additionally, the MSY Bayesian statistics support a male population size reduction (Supplemental Material Figure S5) and very recent coalescence of the majority lineage of the Sila at 1.6 kya (Supplemental Materials Figure S17). Thus, like the Mang, the Sila seem to have independently undergone an extreme bottleneck or founder event. Interestingly, even though these two groups speak languages from different language families and have different genetic profiles, there are strong cultural similarities between the Sila and the Mang in terms of female hair and clothing style, and also in physical appearance (30).
