## SupplementalMaterial Figures for "The paternal and maternal genetic history of Vietnamese populations"

### Supplementary Figures S1-S17

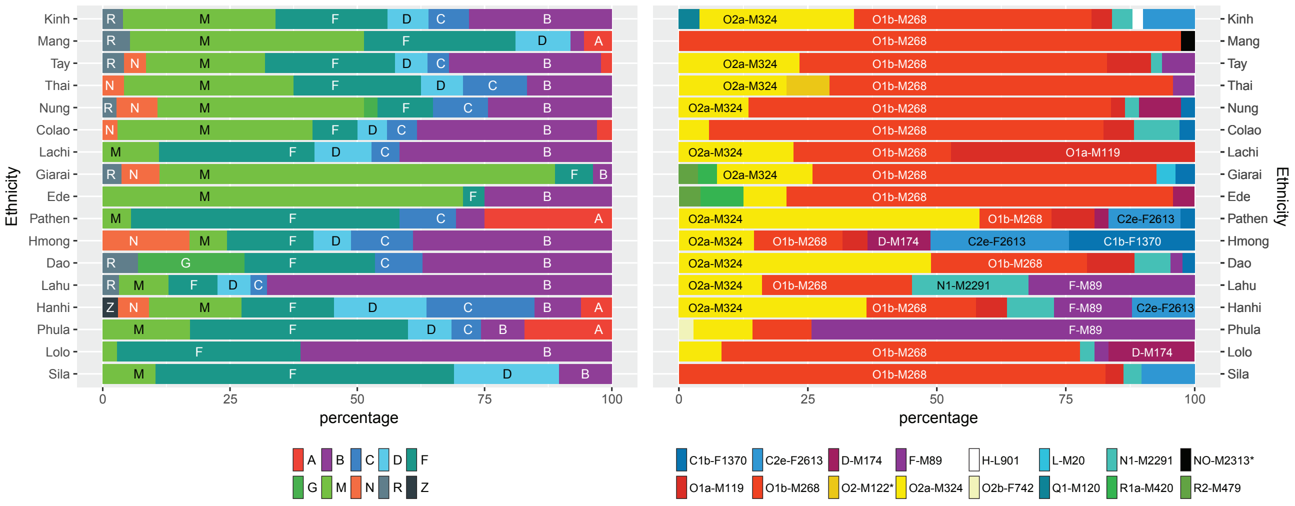


Figure S1 – Macro-haplogroup frequencies per population for mtDNA (left) and MSY (right), in percent.

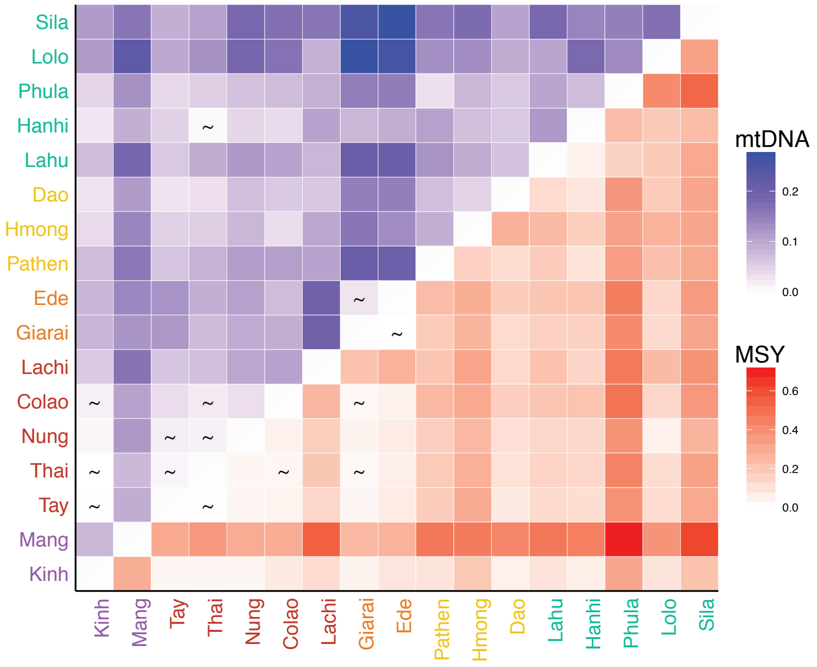


Figure S2 - Pairwise $\Phi_{ST}$ distances between populations, with mtDNA and MSY represented by the blue and red color scale, respectively. Distances that are not significantly different from zero are marked with a tilde symbol ($p > 0.05$). Population labels are color-coded by language family with Austro-Asiatic in purple, Tai-Kadai in red, Austronesian in orange, Hmong-Mien in yellow, and Sino-Tibetan in turquoise.

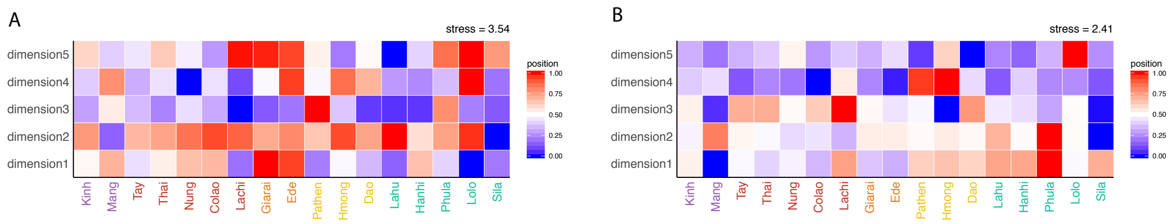


Figure S3 – A: mtDNA and B: MSY multidimensional heatplot based on $\Phi_{ST}$ distances with 5 dimensions and values for each dimension scaled to be between 0 (blue) and 1 (red). Population labels are color-coded by language family with Austro-Asiatic in purple, Tai-Kadai in red, Austronesian in orange, Hmong-Mien in yellow, and Sino-Tibetan in turquoise.

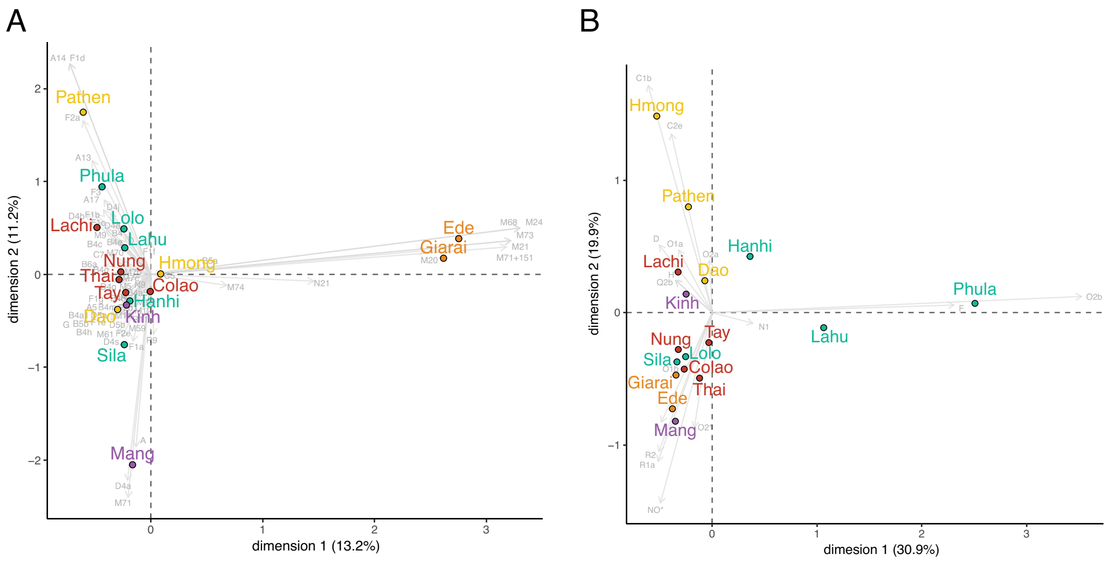


Figure S4 - Correspondence analysis plots for A: mtDNA and B: MSY sequences, based on haplogroup frequencies. Grey vectors and labels show haplogroup projections. Population labels are color-coded by language family with Austro-Asiatic in purple, Tai-Kadai in red, Austronesian in orange, Hmong-Mien in yellow, and Sino-Tibetan in turquoise.

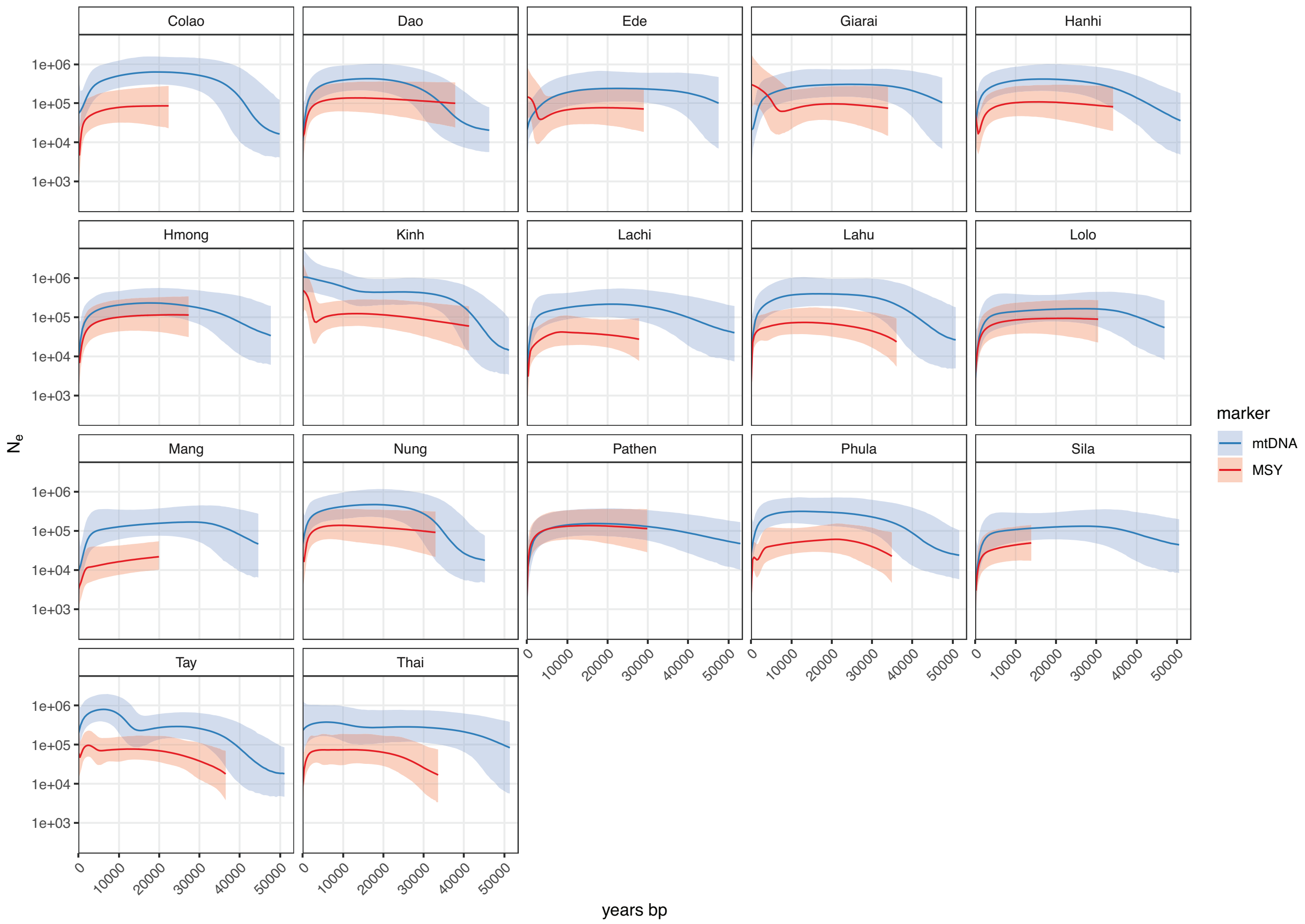


Figure S5 – MSY and mtDNA Bayesian Skyline Plot (BSP) for all populations.

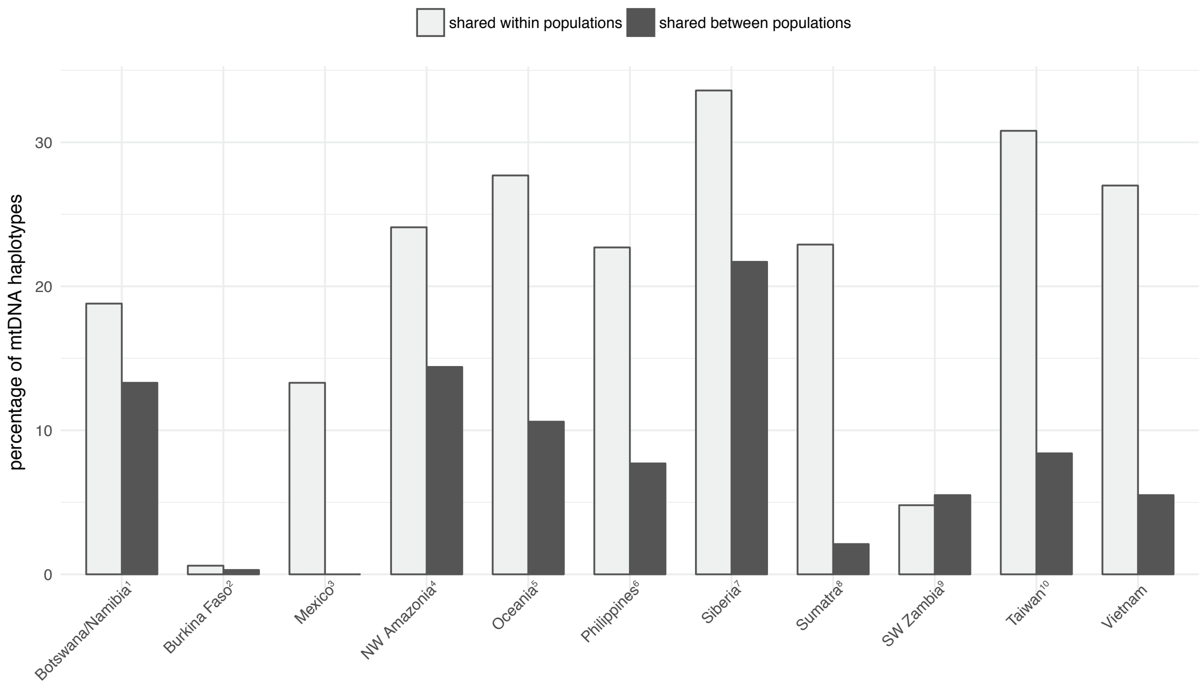


Figure S6 – Proportion of shared mtDNA haplotypes within and between populations in Vietnam vs. other studies of complete mtDNA genome sequences.

1: Barbieri, C. et al. 2014 Am. J. Phys. Anthropol.; 2: Barbieri, C. et al. 2011 Mol. Biol. Evol.; 3: Mizuno F. et al. 2014 J. Hum. Genet.; 4: Arias, L et al. 2018 Am. J. Phys. Anthropol.; 5: Duggan, A. T. et al. 2014 Am. J. Hum. Genet.; 6: Delfin, F. S. et al. 2014 Eur. J. Hum. Genet.; 7: Duggan, A. T. et al. 2013 PLOS ONE. 8: Gunnarsdóttir, E. D. et al. 2011 Nat. Commun.; 9: Barbieri, C. et al. 2013 Am. J. Hum. Genet.; 10: Ko, A.-M.-S. et al. 2014 Am. J. Hum. Genet.; Vietnam: this study.

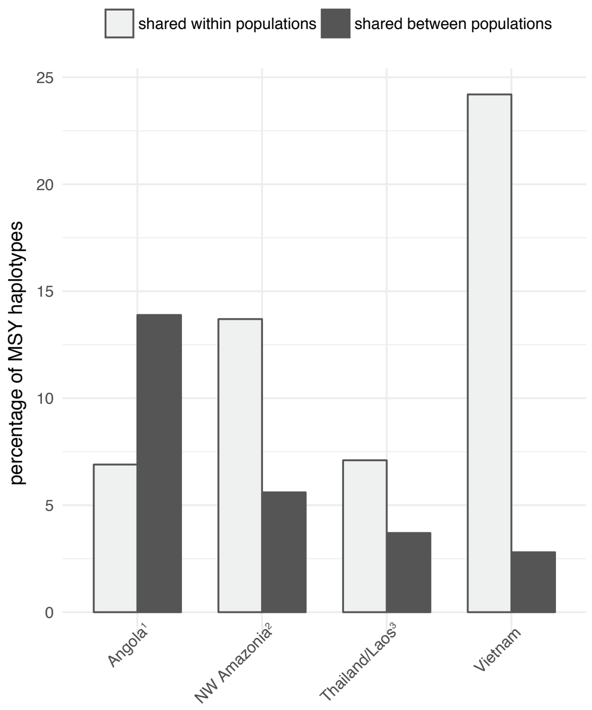


Figure S7 - Proportion of shared MSY haplotypes within and between populations in Vietnam vs. other studies of the same MSY region.
1: Oliveria et al. 2019 Hum. Genet; 2: Arias et al. 2018 Am. J. Phys. Anthropol.; 3: Kutanan et al. 2019 Mol. Biol. Evol.; Vietnam: this study.

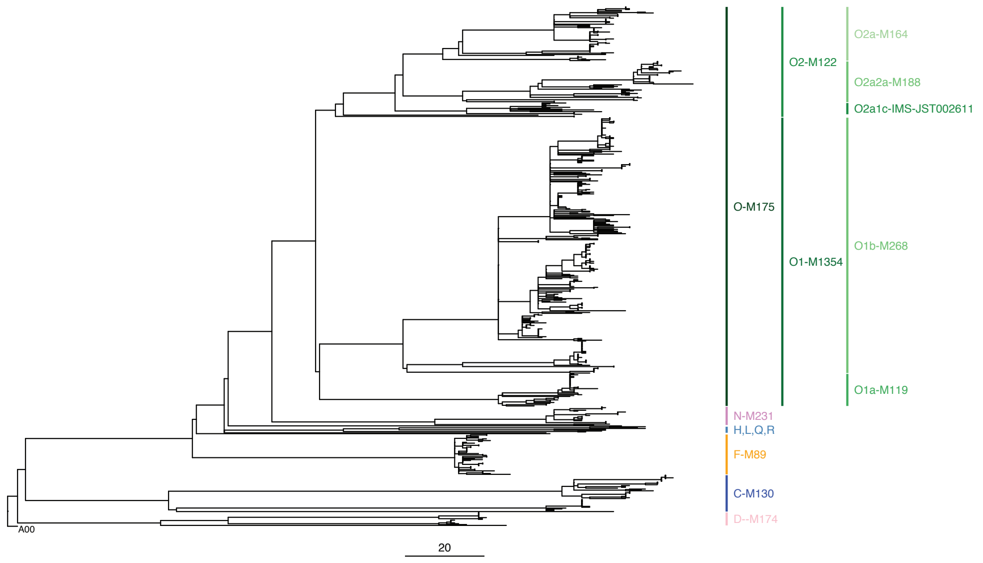


Figure S8 – Maximum parsimony tree of Vietnamese MSY sequences labeled with macro-haplogroups.

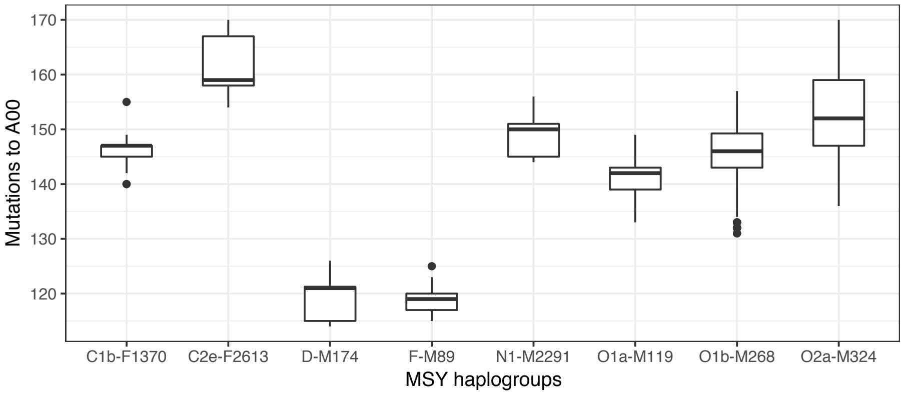


Figure S9 – Comparison of the number of mutations to A00 for selected MSY macro-haplogroups.

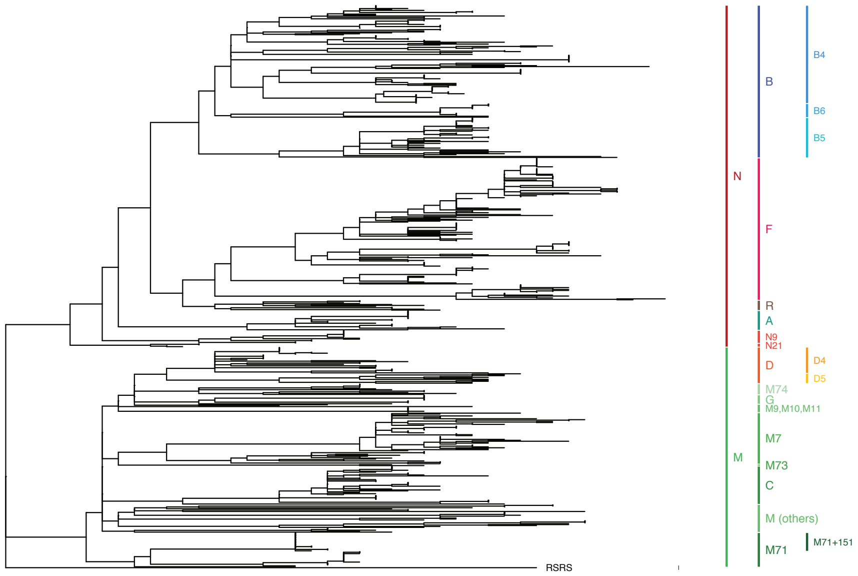


Figure S10 – Maximum parsimony tree of Vietnamese mtDNA sequences with haplogroup labels.

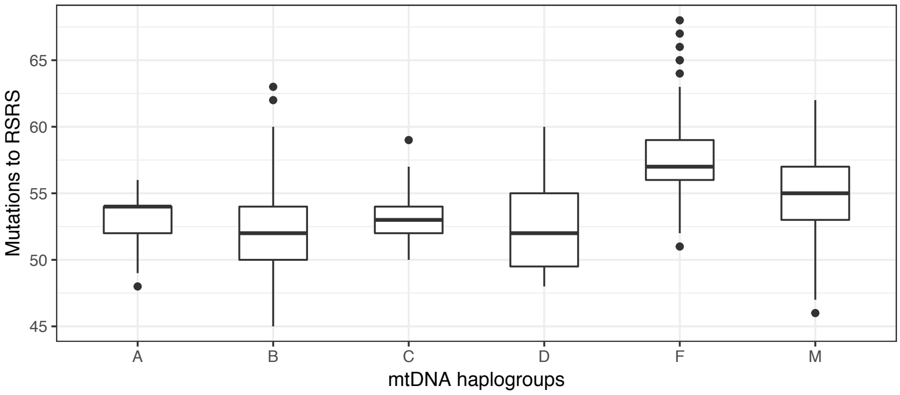


Figure S11 – Comparison of the number of mutations to the RSRS for selected mtDNA macro-haplogroups.

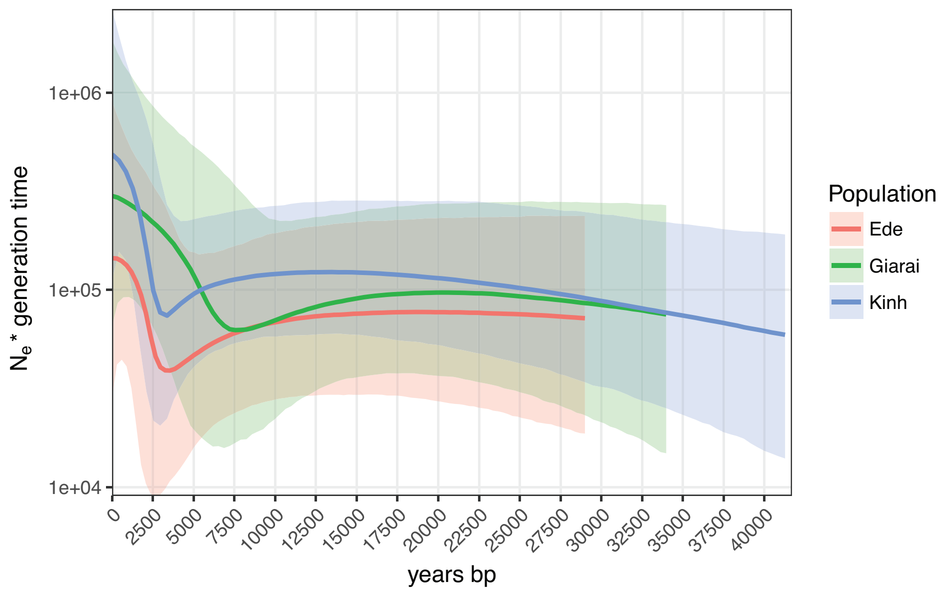


Figure S12 - MSY Bayesian Skyline plot (BSP) for the Giarai, Ede, and Kinh.

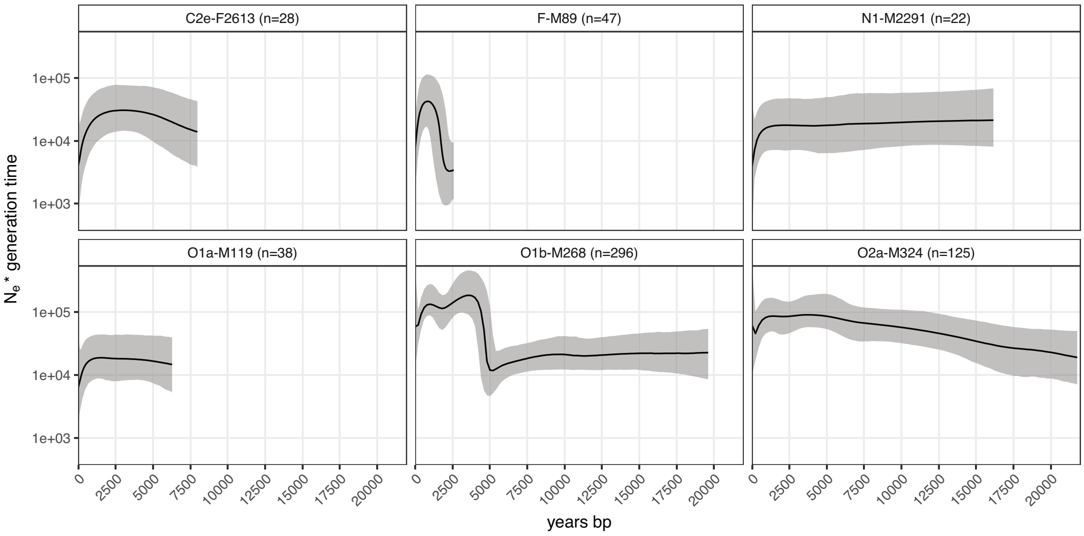


Figure S13 – BSP of major MSY
macro-haplogroups, including sub-groups with a sample size > 20: C2e-F2613, F-M89, N1-M2291, O1a-M119, O1b-M268, and O2a-M324.

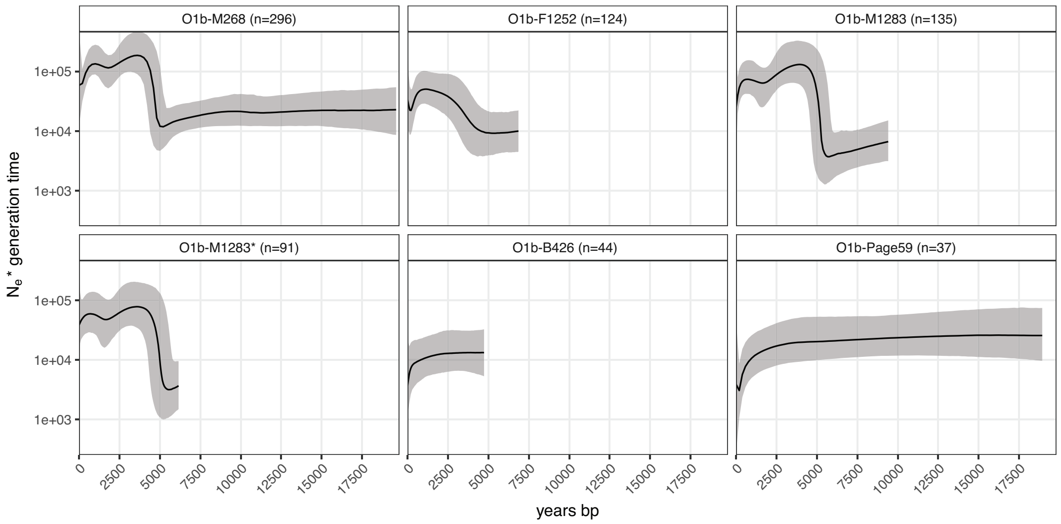


Figure S14 – BSP of MSY macro-haplogroup O1b-M268 sub-haplogroups. The asterisk denotes a haplogroup without subgroups.

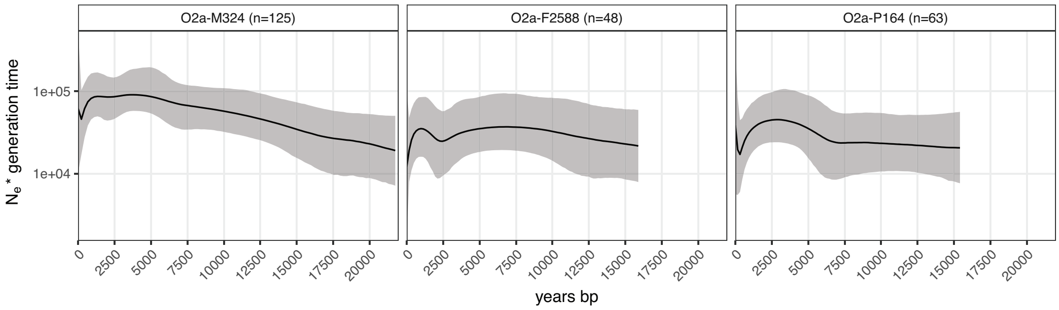


Figure S15 – BSP of MSY macro-haplogroup O2a-M324 and sub-haplogroups.

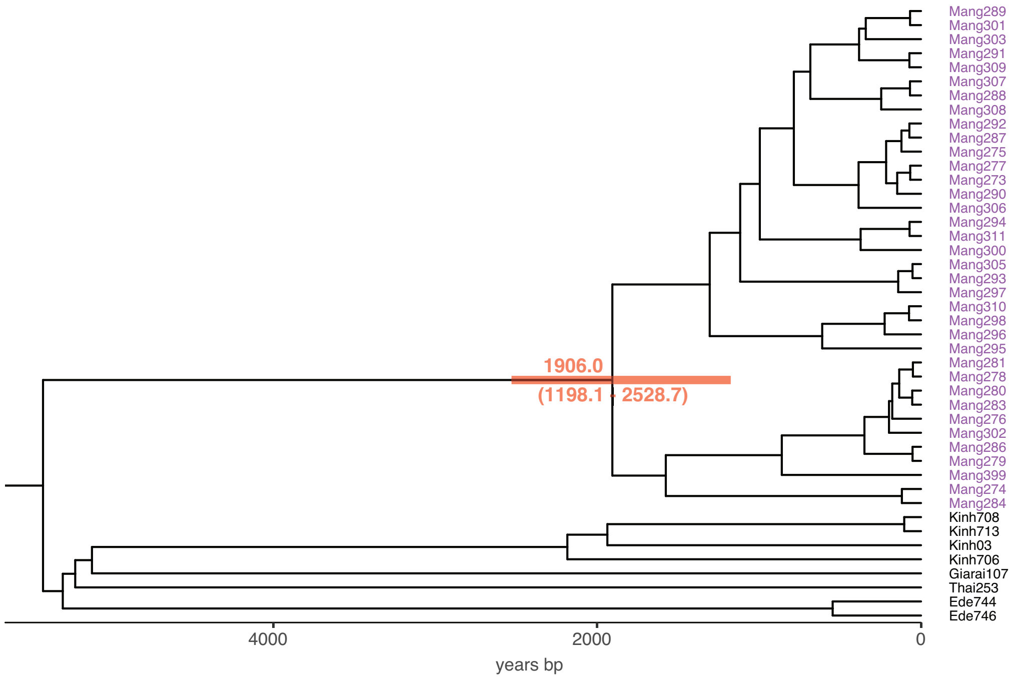


Figure S16 – Bayesian tree of MSY haplogroup O1b-B426 (O1b1a1a1b1a) including the cluster of Mang individuals. The TMRCA point estimate and 95% HPD interval are shown in red.

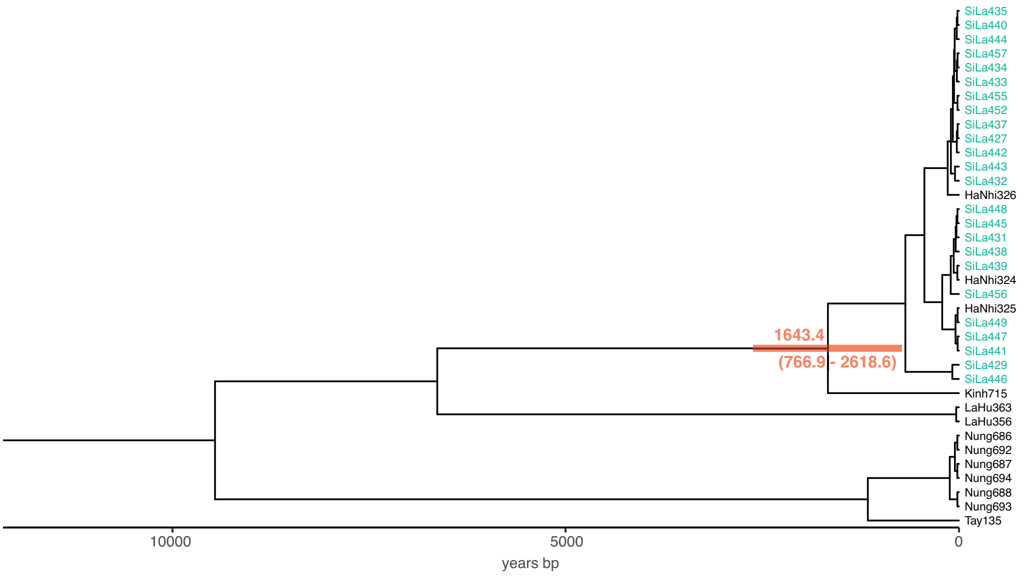


Figure S17 – Bayesian tree of MSY haplogroup O1b-F1759 (O1b1a2a1) including the cluster of Sila individuals. The TMRCA point estimate and 95% HPD interval are shown in red.
